# The quantity of Neanderthal DNA in modern humans: a reanalysis relaxing the assumption of constant mutation rate

**DOI:** 10.1101/065359

**Authors:** William Amos

## Abstract

Few now dispute that a few percent of the DNA of non-African humans is a legacy of interbreeding with Neanderthals. However, heterozygosity and mutation rate appear to be linked such that the loss of diversity associated with humans migrating out of Africa caused a mutational slowdown, allowing Africans to diverge more from both our common ancestor and Neanderthals. Here I use a range of contrasting tests aimed at distinguishing between mutation slowdown and introgression as explanations for the higher rates of base-sharing between non-Africans and Neanderthals. In every instance the mutation slowdown hypothesis fits better. Thus, while some interbreeding likely occurred, as evidenced by the finding of skeletons of admixed individuals, adaptive genes and the apparently large contribution of Denisovan DNA to Oceanian genomes, my results challenge the idea that non-Africans generally carry an appreciable Neanderthal legacy. My analysis shows that inferences about introgression may be unreliable unless variation in mutation rate linked to demographically induced changes in heterozygosity can be excluded as an alternative hypothesis.

## Introduction

It is widely accepted that modern non-African humans carry a few percent of Neanderthal DNA as a legacy of historical inter-breeding ^1-3^. The original inferences were based on higher than expected levels of base-sharing between Neanderthals and non-Africans. This pattern was subsequently reinforced by analyses of the size of introgressed fragments ^4^, analysis of the Denisovan genomes ^5^ and targeted analyses of specific individuals ^6^ and genes ^7^. The latter two are convincing but inferences of the more general patterns explicitly assume that mutation rate is constant ^1,8^. In fact, human mutation rate likely varies between populations ^9^ and this variation covaries with heterozygosity (heterozygote instability, HI) ^10^, a pattern also reported recently in plants ^11^. Consequently, the large loss of heterozygosity that humans suffered while migrating out of Africa lowered the mutation rate in non-Africans ^10^, causing non-Africans to be more related than Africans both to our common ancestor and to related lineages like Neanderthals. In view of this I decided to conduct a series of tests aimed at distinguishing between Neanderthal introgression and mutation slowdown.

Inter-breeding was originally inferred through the so-called ‘ABBA-BABA’ test, also referred to as D statistics ^12^, which quantifies asymmetrical base sharing in 4-way DNA sequence alignments, here two humans, the Neanderthal and the chimpanzee (H1-H2-N-C). Bases of interest are those where the two modern humans differ and where one base matches the chimpanzee while the other matches the Neanderthal (‘ABBA’ and ‘BABA’). Under a null model, ABBA and BABA sites should occur at equal frequencies. That the counts are significantly asymmetrical was used to infer introgression, but the asymmetry is also explicable by a greater mutation rate in Africans (**Fig 1**) if the back-mutation rate is sufficiently high.

**Figure 1.**
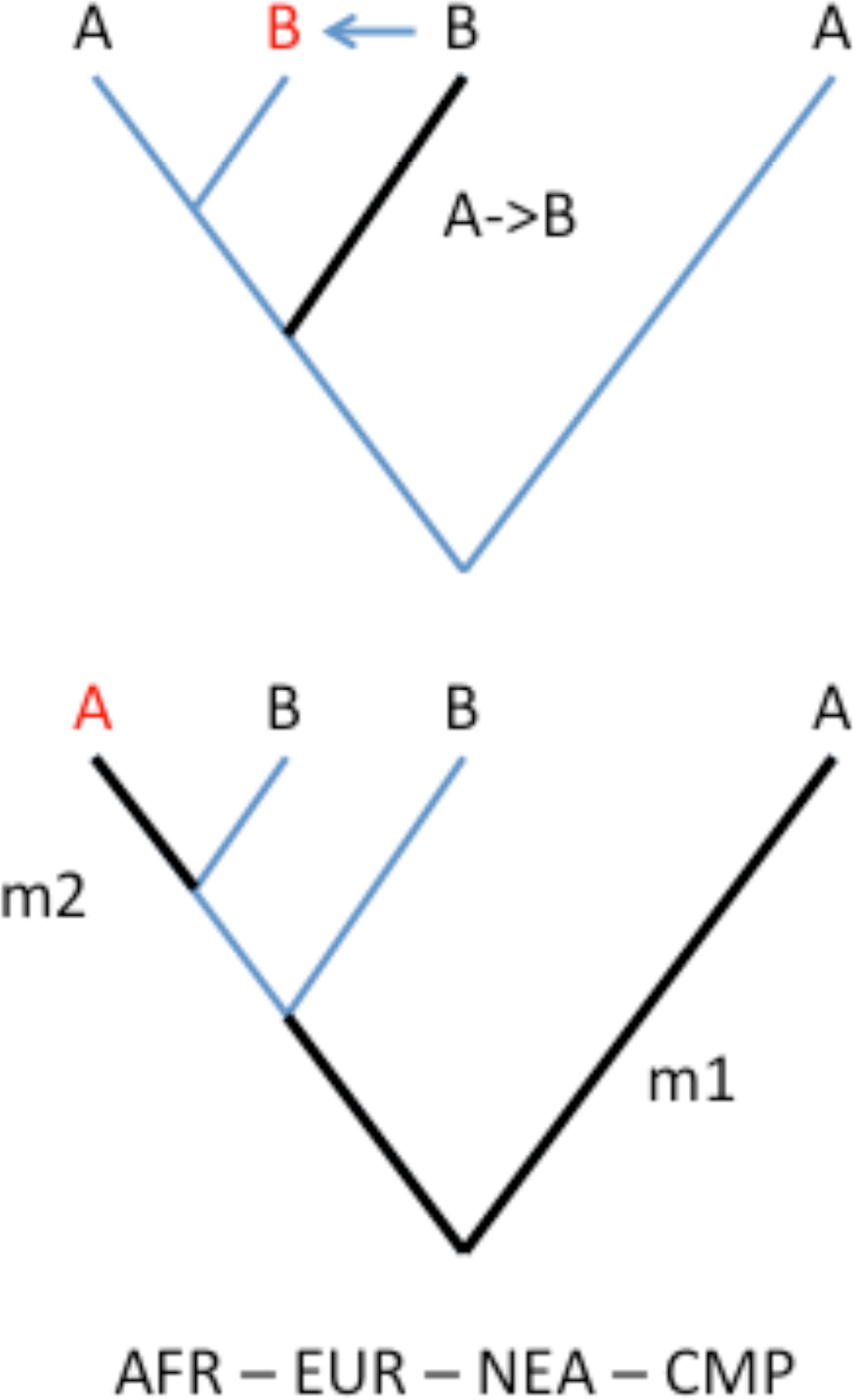
Alternative hypotheses to explain excess base sharing with Neanderthals. Each tree depicts a mechanism by which an “ABBA” pattern is generated, wherein a European human (EUR) shares a base with the Neanderthal (NEA) while, at the same site, an African human (AFR) shares a base with the chimpanzee, *Pan troglodytes* (CMP). Both trees assume only two states, ‘A’ and ‘B’. Heavy black lines indicate lineages on which mutations occurred. In the top tree, a single mutation creates a new allele in NEA. This new allele, ‘B’, then enters EUR by introgression. In the lower panel a first mutation (m1) between the chimpanzee and the human – Neanderthal ancestor creates a “BBBA” pattern. A second (back) mutation (m2) then recreates an ‘A’ allele in AFR. For calculations relating to the required rate of back mutations see **Supplementary material 1**.

### Testing whether back-mutations provide a plausible alternative explanation

If mutations are distributed evenly and randomly across the genome, too few back-mutations are expected genome-wide (**Supplementary Material 1**). However, mutation rate varies hugely between genomic regions and sites ^13 14^. Since the probability of a back-mutation scales with mutation rate squared, a high variance in mutation rate could increase greatly the expected number of back-mutations. To illustrate, if all mutations occur at just 10% of sites (i.e. 90% of sites are invariant), the number of back-mutations would rise ten-fold. The back-mutation rate would rise further wherever mutations are clustered ^15-17^ and further still if heterozygous sites attract gene conversion events that ‘correct’ the heterozygous site itself ^18^.

The idea that back-mutations occur sufficiently frequently to be considered as a viable alternative hypothesis to introgression is supported empirically by base combination counts in Green et al. for a San-French-Neanderthal-chimpanzee alignment (their Supplementary Material p138) ^1^. CBBA and BCBA both require two mutations including at least one transversion, and both occur over 6,000 times in ~25% of the genome. Since the probability of a transition is approximately four times that of a transversion, these data imply ~96,000 instances each of ABBA and BABA generated by back-mutations genome-wide. This figure is consistent with a direct counting approach (see Supplementary Materials 1) and is of the correct order required for the observed ABBA-BABA asymmetry to be explained by a model based on mutational slowdown.

### The signal of introgression is independent of Neanderthal sequences

Mutation slowdown predicts that the ABBA-BABA signal is an inherent property of modern humans, arising through a higher African mutation rate. Such a signal will be present whether or not the Neanderthal genome is present. I therefore conducted an ABBA-BABA analysis, substituting the Neanderthal genome with ancestral human alleles, AA, as inferred from a panel of primates by the 1000 genomes project, i.e. H1-H2-AA-C. To prevent any possible spill-over signal, all informative Neanderthal sites that contributed to the original ABBA-BABA signal were excluded. The resulting signal is even stronger than for Neanderthals (**Fig 2**). The presence of a signal that not only remains but actually increases in strength when Neanderthal information is excluded indicates that much or all of the signal is unrelated to Neanderthal introgression and therefore requires an alternative explanation **(Supplementary material 2**).

**Figure 2.**
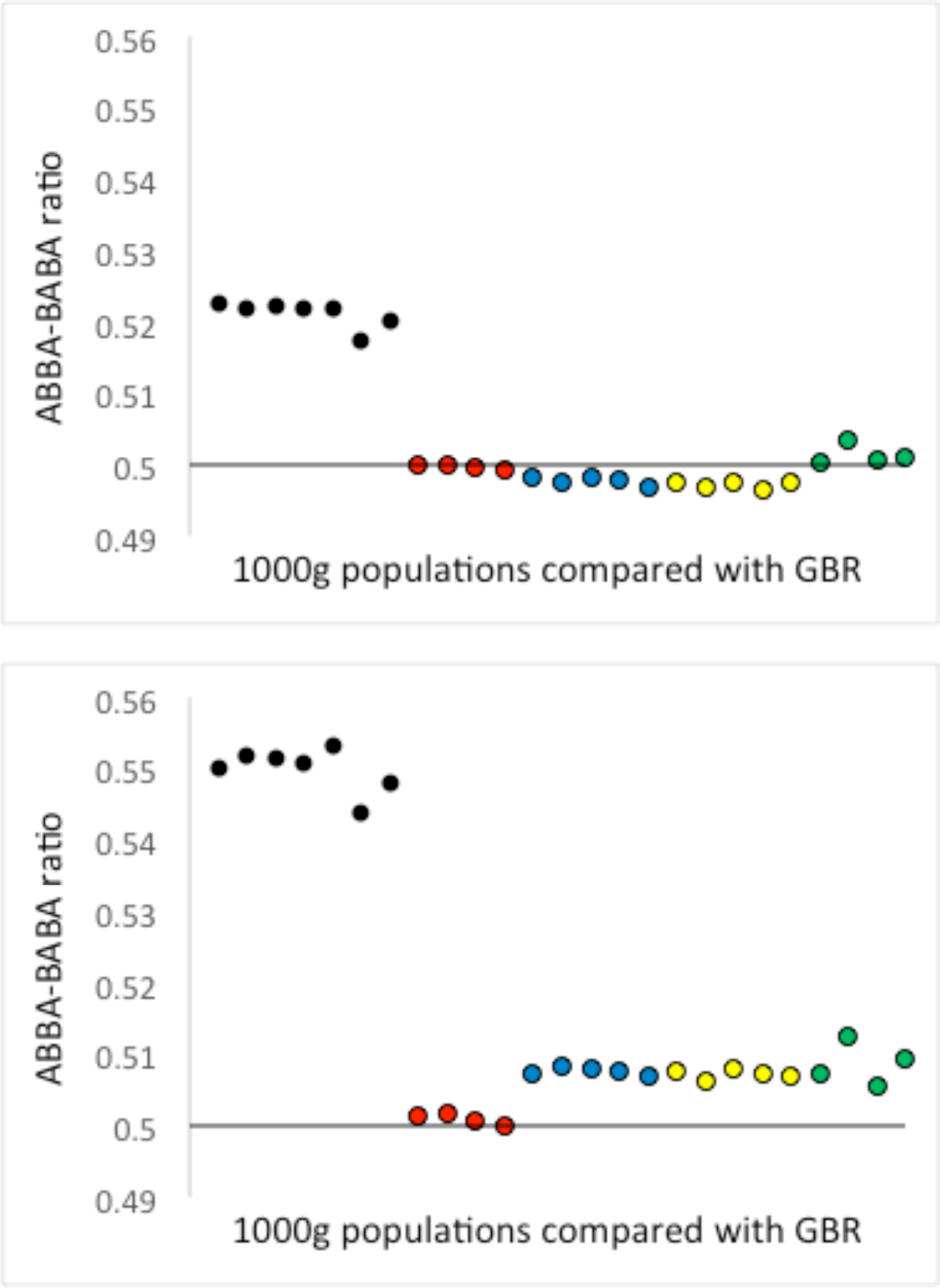
Testing for introgression without the Neanderthal genome. Each dot represents an ABBA-BABA test of the form X-GBR-Test-Chimpanzee, CMP, where X is one of 25 populations in the 1000 genome Phase 3 data and Test is the taxon being tested for evidence of introgression. Human populations are colour-coded to indicate geographic region: black=Africa; red=Europe; blue=southern Asia; yellow=East Asia; green=America, populations appearing in the same order as listed in methods. Standard errors of the mean are of the order 0.001 and are too small to show. The results are expressed on the Y axis as the proportion of all autosomal counts of ABBAs and BABAs that are ABBA, where ABBA is defined as the state where the test taxon shares a base with GBR. Thus, values above 0.5 would previously have been interpreted as evidence of introgression into GBR. In the top panel the test taxon = Neanderthal, NEA. The values for Africa, around 0.52, equate to an excess of around 8% ABBAs and are essentially identical to the results obtained by Green et al. In the bottom panel Test = Ancestral Allele, AA, *sensu* the 1000 genomes project. All informative sites contributing to the signal found in the top panel for X-GBR-NEA-CMP were excluded from the X-GBR-AA-CMP analysis.

### Modern human mutation rate is higher in Africa than outside

The base combinations listed by Green et al. (their Supp. Mat. P. 138) offer further insight ^1^. Just like ABBA and BABA, base combinations where only the two humans differ, BAAA and ABAA, are asymmetrical to about the same degree, with counts of 756324 and 689594. Taken at face value, this asymmetry indicates 10% more substitutions on the African lineage, as reported previously^10^. Despite the large number of bases involved and the use of high coverage genomes, the BAAA:ABAA asymmetry is ascribed entirely to sequencing errors. To clarify the underlying cause I examined the distribution of derived alleles Since most rare alleles are recent ^19^, alleles with frequencies under 10% likely arose post ‘out of Africa’. After excluding singleton variants, any given rare derived allele is 3.37 times as likely to be found in an African compared with a non-African (**Supplementary material 3**), a strong pattern confirming that mutation rate is higher in Africa.

### Discordant fragments in non-Africans are very rare

If present, introgressed archaic fragments should be detectable directly, without relying on alignments to the Neanderthal genome. On average, sequence divergence is greater between Neanderthals and any modern human than between any two modern humans. I therefore calculated all-against-all intrapopulation pairwise divergences for non-overlapping 20Kb windows across the genome using data from the 1000 genomes project ^20^. In most windows, maximum within-population pairwise divergence (MWD) is higher inside Africa than outside, consistent with the higher diversity found in Africa ^21^ (**Fig 3a**). Moreover, the few instances where MWD(non-African) is appreciably greater than MWD(Africa) rarely coincide with inferred peaks of Neanderthal introgression (**Fig 3b**). Importantly, the frequency and magnitude of such windows on the X chromosome, where introgression is thought to be minimal or absent, tend to be greater than equivalent autosomal values (**Supplementary Material 4**).

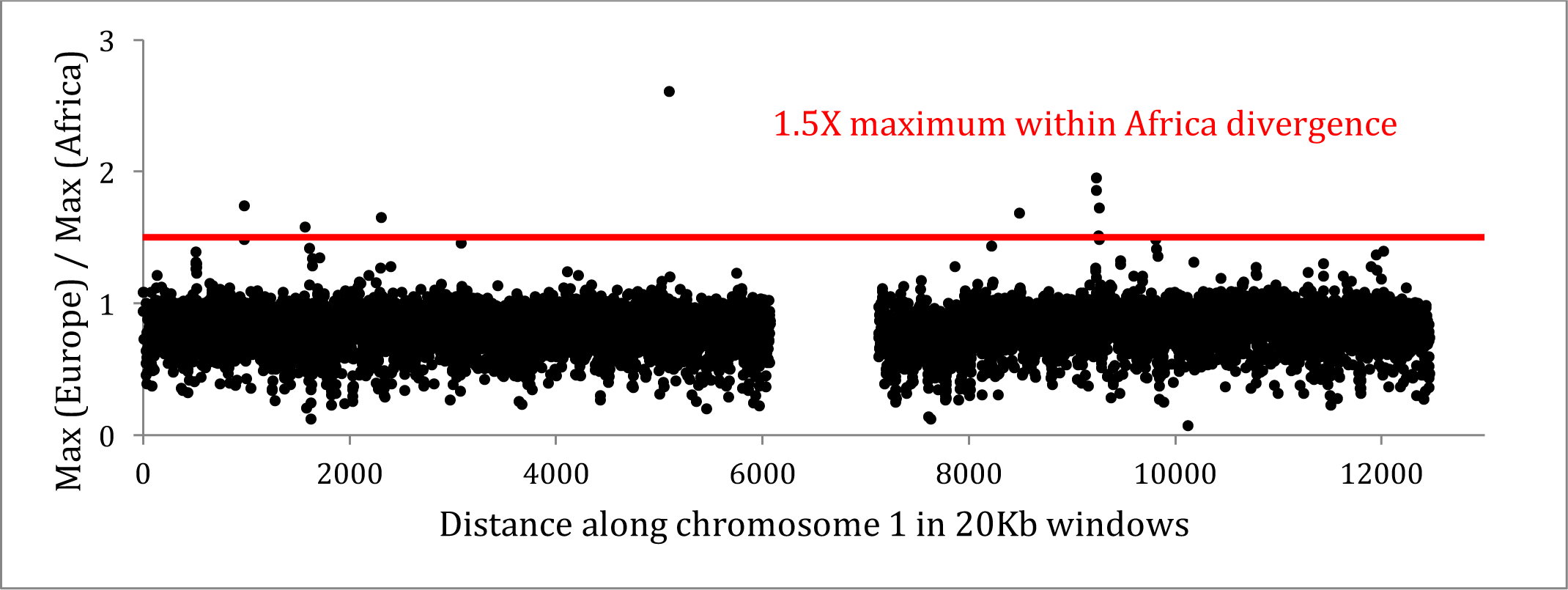

**Figure 3.**
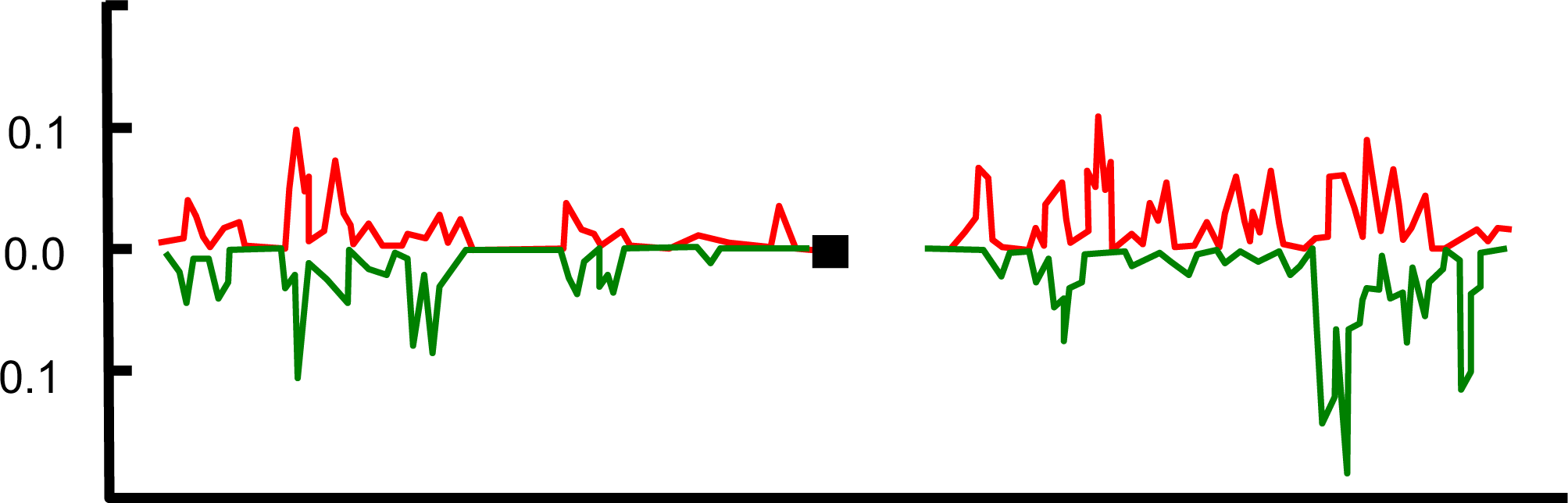
Direct test for presence of unusual introgressed fragments. Figure 3a shows statistics based on 1000 genome Phase 3 data for human chromosome 1, partitioned by non-overlapping 20Kb window. Within each window, all possible within population inter-individual distances are calculated by counting the number of base differences. The maximum values found in the five African-origin populations (LWK, ESN, MDL, GWD, YRI, = MAX(Africa)) and five European (GBR, FIN, CEU, TSI, IBS = MAX(Europe)) were recorded, yielding a ratio MAX(Europe) / MAX(Africa), on the Y-axis. The overwhelming majority of windows give ratios less than 1.5 and average ~0.8, reflecting the greater diversity in Africa. The highest ratio observed here is 2.68, well short of the average expected value for Neanderthal DNA. Figure 3b is redrawn from Sankararaman et al.^22^ and gives the map of inferred Neanderthal ancestry in red (Europe) and green (East Asia), expressed as ‘proportion of confidently inferred Neanderthal alleles’. All other chromosomes given similar patterns (data not shown). Note that there is little or no correspondence between peaks in Figures 3a and 3b.

### Non-Africans carry less than 0.2% divergent DNA fragments

To estimate the archaic fraction in individual non-African genomes I used a parallel approach. I assume that most introgressed fragments are neutral and, having been introduced around 2,000 generations ago, are unlikely to have reached high frequency (>50%) within a population. Since some unknown proportion of genomic windows will show too little human-archaic divergence for the archaic sequence to stand out, I ‘spiked’ each non-African population with a randomly selected African individual. I assume that the number of windows where this African individual qualifies as being unusual sets a conservative minimum for the number of windows in which an archaic sequence would be detected using the same criteria **(Supplementary Material 4**). This approach reveals that, on average, 0.14% of a typical non-African genome can be considered unusually divergent, an order of magnitude less than published estimates. The true value will probably be lower still because my methods are conservative. In addition, some unusually divergent sequences will arise through chance and / or selection rather than introgression.

### Exploring the relationship between Neanderthal-human base-sharing and heterozygosity

Mutation and introgression can also be distinguished by their relationship with heterozygosity. Introgression predicts the ABBA-BABA signal will be either uncorrelated or negatively correlated with the difference in heterozygosity between populations (*H_diff*), depending on whether introgressed fragments appreciably increase heterozygosity in the recipient population. Mutation slowdown predicts the exact opposite. Under this hypothesis the ABBA-BABA signal will be strongest wherever the difference in mutation rate is greatest, and under HI this is strongly correlated with *H_diff*^10^. Mutation slowdown therefore predicts a strong positive correlation. Plotting *H_diff* against the ABBA minus BABA difference for 100Kb windows across the genome reveals a strong positive correlation in all pairwise population combinations, exemplified one European-African comparison and one European-East Asian comparison (**Fig 4**). These correlations are difficult to reconcile with a model where the ABBA-BABA signal is driven largely or wholly by introgression and it is difficult to understand why or how relatively high heterozygosity in Africa would increase the chance of an introgressed fragment being retained. Conversely, these correlations are expected, indeed required by a model based on heterozygosity-driven mutation slowdown.

**Figure 4.**
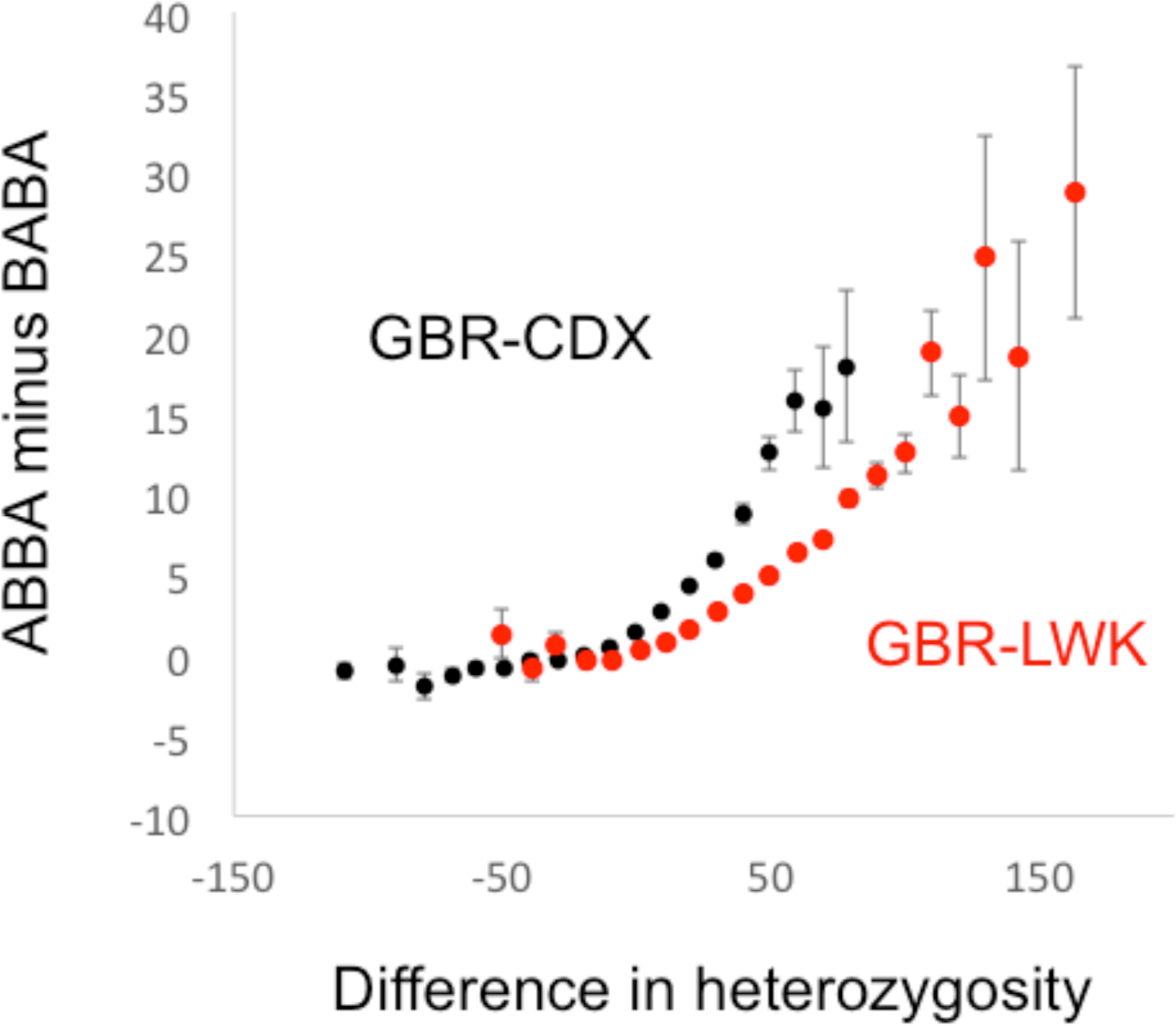
Difference in heterozygosity between modern human populations predicts ABBA-BABA signal. The genome was divided into non-overlapping 100Kb windows. Within each window, the expected number of heterozygous sites in each population was calculated as 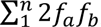, *n* sites, *f*_*a*_ and *f*_*b*_ the frequencies of the two alleles, yielding a value for heterozygosity difference (X-axis). In each window I also calculated the difference in expected number of ABBA and BABA sites, assuming Hardy-Weinberg equilibrium (y-axis, see Methods). The two population pairs presented are GBRLWK and GBR-CHB. All other GBR-Africa comparisons yield profiles that are essentially identical to GBR-LWK. Similarly, all other GBR-East Asia comparisons yield profiles that are essentially identical to GBR-CHB. The strong, positive non-linear correlations are not explicable by artefact and support the mutation rate model. Under the introgression model the observed correlation should be absent or negative. The different trajectories reflect the different timescales over which heterozygosity changes and new mutations accumulate.

### Which alleles are rare?

The two hypotheses also give opposing predictions for the populations in which the ‘A’ and the ‘B’ alleles are rare. Both hypotheses involve the relatively recent introduction of largely neutral alleles into modern humans, and such alleles will tend to be rare. Under introgression, we expect the Neanderthal ‘B’ allele to be rare outside Africa. Conversely, under mutation slowdown, it will be the ‘A’ allele that will be rare in Africa, implying that the ‘B’ is common. To test these predictions, I partitioned the ABBA-BABA counts for X-GBR-NEA-CMP alignments according to *f*, the frequency of the ‘B’ allele in GBR. Almost no signal is present when *f* is at intermediate frequency (10% > *f*< 90%). However, the signal switches between polar opposites depending on whether ‘B’ is rare (*f* < 10%) or common (*f* > 90%). The original ABBA-BABA signal used to infer introgression corresponds to the pattern seen when ‘B’ is common and ‘A’ is rare, and hence appears to be driven by mutation slowdown.

**Figure 5.**
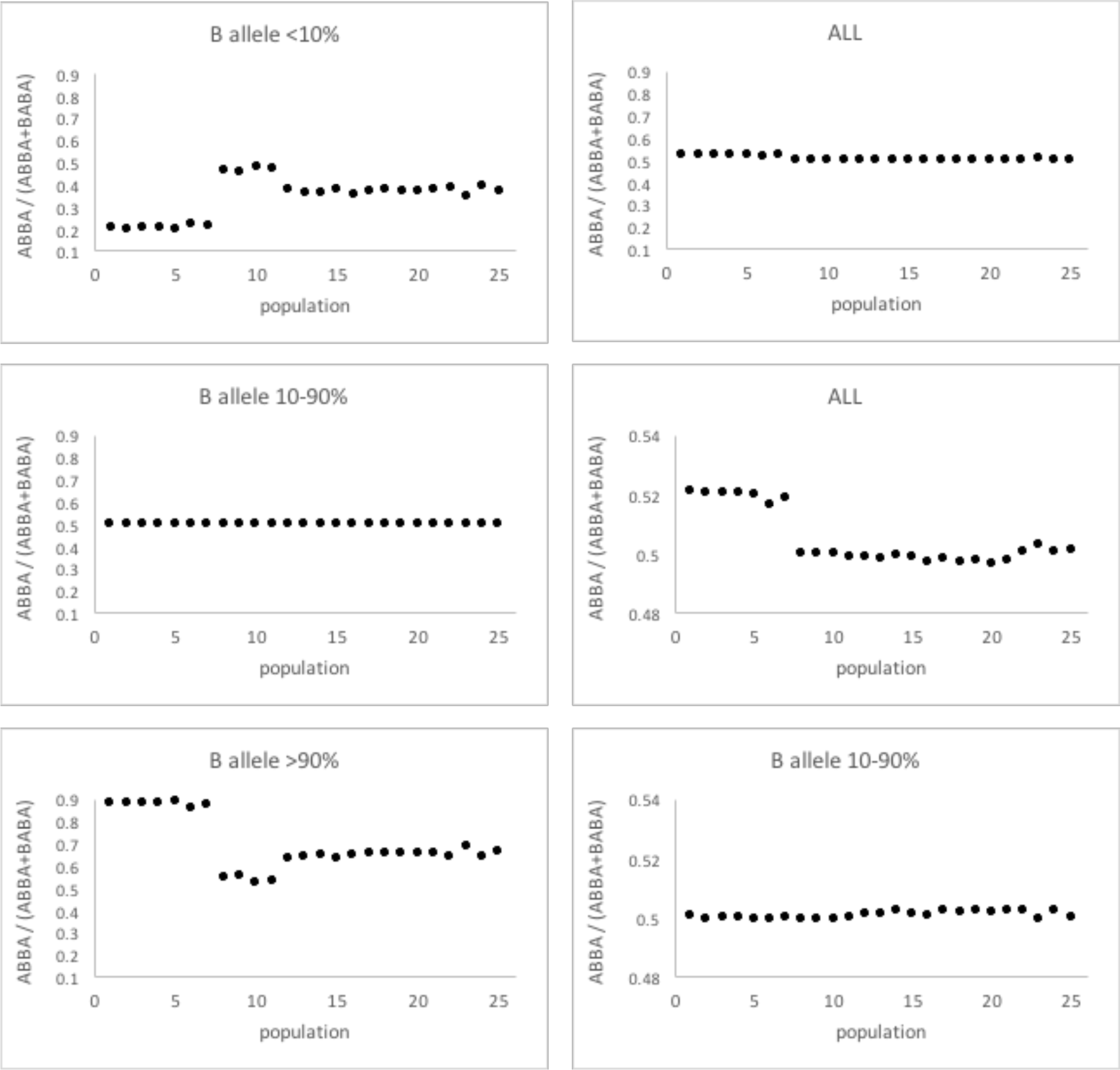
Dependence of the ABBA-BABA signal on the frequency of the ‘B’ allele in Europe. A standard ABBA-BABA analysis was conducted based on the tetrad X-GBR-NEA-CMP with ABBA and BABA counts stored according to the frequency of the ‘B’ (i.e. Neanderthal) allele in Europe (GRB). Axes are the same as in Figure 2, with populations appearing in the same order as listed in methods. Panels on the lefthand side show, from top to bottom, ABBA-BABA ratios for ‘B’ at low, intermediate and high frequency. The original ABBA-BABA signal reported by Green et al. is show top right (same scale as the lefthand panels) and middle (expanded Y-axis scale). The entire signal is driven by extreme frequency alleles, as seen in the bottom right panel that show a lack of detectable signal when the ‘B’ allele is at intermediate frequency, even when the Y-axis scale is expanded. Note, the original signal (centre right panel) equates to the case when ‘B’ alleles are common in GBR, the exact opposite to the expectation under introgression.

### What is the source of human ABBA-BABA signal variation?

As a final test I looked at the relative impact of varying which human populations are included in the analysis. Under the introgression hypothesis, Africans are seen a passive controls that carry little or no introgressed material, though see also ^23^. Consequently, most variation in the strength of the ABBA-BABA signal under introgression will derive from variation in Neanderthal content of the non-African populations. Conversely, under mutation slowdown, non-African populations are seen as being rather homogeneous such that most variation in the ABBA-BABA signal will be associated with variation in mutation rate among the African populations. To test which of these two scenarios fits best I conducted ABBA-BABA tests of the form X-Y-NEA-CMP, where X is represents one of five African populations (LWK, ESN, MSL, ASW, YRI) and Y is any one of the five European populations. Each of the 10 populations thus has five ABBA-BABA comparisons from which a variance is calculated. The ABBA-BABA scores vary significantly more when a given European population is compared with different African populations than *vice versa*. As elsewhere, this patterns appears more consistent with mutation slowdown than with introgression.

**Figure 6.**
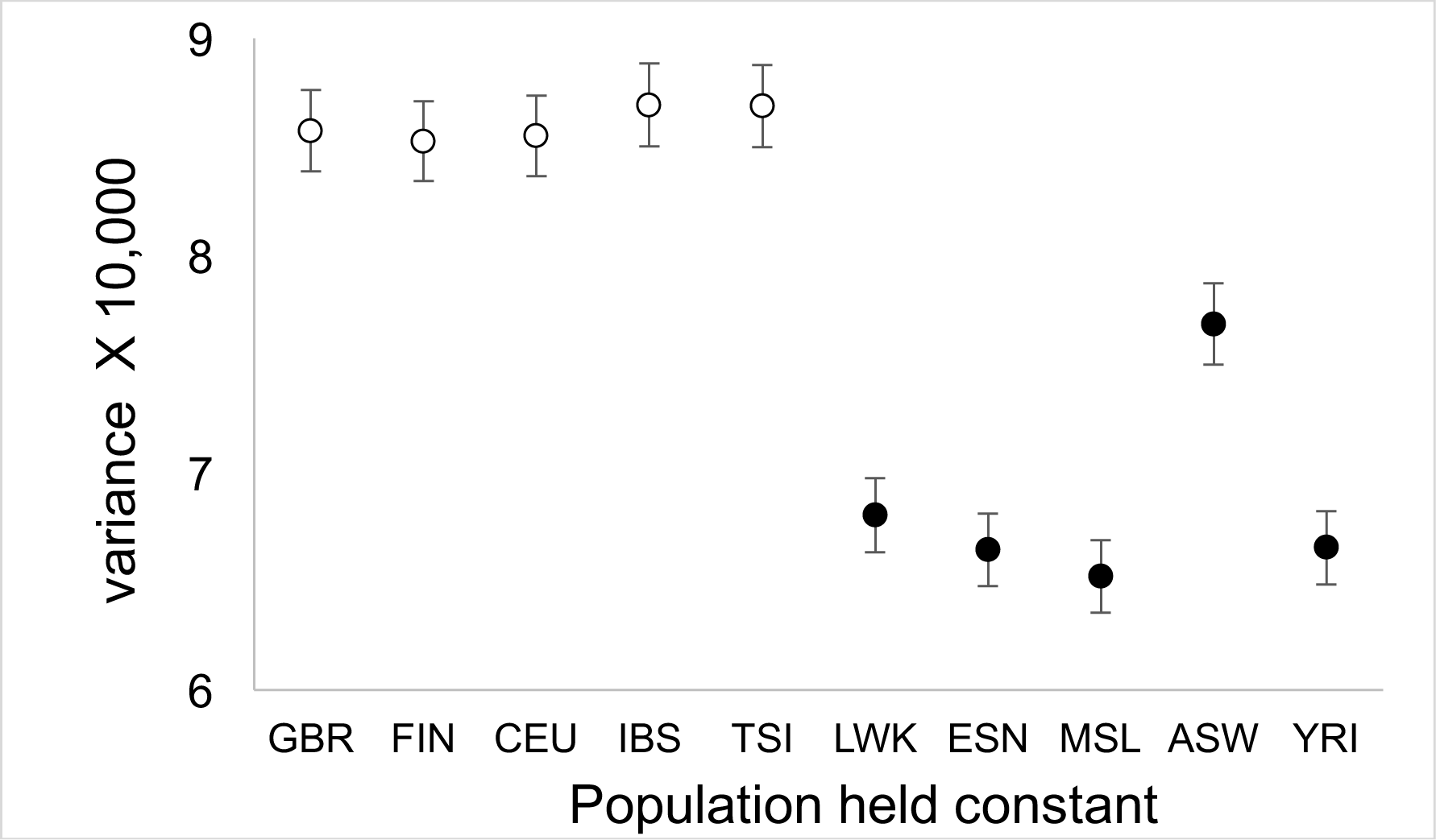
The main source of variation in ABBA-BABA signal. To test whether African or non-African populations are the main source of variation in the ABBA-BABA signal the autosomal genome was divided into non-overlapping 100Kb blocks. Within each block, all pairwise African – non-African comparisons were conducted between five European populations (GBR, FIN, CEU, IBS, TSI) and five African populations (LWK, ESN, MDL, ASW, YRI), yielding 25 ABBA-BABA scores. Each population thus has five comparisons over which variance is calculated. Variance in ABBA-BABA signal is consistently greater when the European population is held constant and the African population is varied than when the African population is held constant and the European population varied. Note how the admixed African population, ASW, is intermediate.

### Discussion

Previous studies have made a compelling case for a Neanderthal legacy in modern non-Africans. However, until now, the only alternative hypothesis was based on historical population sub-structure ^24^, a possibility that is debated ^25^ and difficult to resolve directly. Mutation slowdown provides a completely new alternative. Crucially, the mutation slowdown model makes a range of predictions that can be used to help distinguish it from introgression. Perhaps unexpectedly, in every test I applied the mutation slowdown model offers a better, often unambiguously better, fit to the data than the Neanderthal introgression model. My results therefore appear to be at odds with previous studies, a conflict that can be resolved in three main ways.

First, my analyses are constrained both by the populations covered by the 1000 genomes project, which do not include representatives from Oceania, and refer mainly to broad patterns linked to Neanderthals. As such, they have no particular implications either for the Denisovan story ^26^, nor for anecdotal observations of specific individuals or genes. Indeed, evidence for occasional interbreeding events seems rather strong and, if the resulting offspring survived and bred, this may well have led to the selective retention of particular alleles that proved beneficial in a modern background.

Second, much of the strongest evidence for introgression is based on higher order patterns. For example, the inferred size of introgressed blocks is largest in archaic modern humans and gets progressively smaller towards the present ^4^, as expected if introgressed fragments are progressively broken down by recombination. These patterns appear convincing but, like the original ABBA-BABA analysis, have yet to be challenged seriously by an alternative model. As with ABBA-BABA, mutation slowdown now offers a possible alternative. Key is the possibility that recombination rate and mutation rate appear correlated ^27, 28^, plausibly because a proportion of the gene conversion events invoked by HI will be resolved by crossing over.

With mutation rate and recombination rate correlated, the ‘out of Africa’ bottleneck would have caused both to reduce in tandem. The result would be large blocks, corresponding to regions where mutation rate and recombination differ most between Africans and non-Africans, patterns that would like be accentuated by non-independence ^29,17^. Just as if these blocks had arisen by introgression they would subsequently break down by recombination, a process that would be accelerated by the recovery of both mutation and recombination rates. Moreover, the original loss of heterozygosity was modulated by natural selection, particularly at immune-related genes, providing a potential link between signals interpreted as introgression and functional genomic regions.

In terms of explaining variation in block size, the HI model remains speculative and difficult to test due to lack of knowledge about key parameters. However, it gains considerable credence from the analysis presented here, where all aspects of the ABBA-BABA test I looked at seem better explained by mutation slowdown. Consequently, while it would be premature to argue that the higher order analyses of aspects such as block size can be discounted, I believe enough doubt has been cast to warrant revisiting. In particular, it seems important to allow for the possibility that mutation rate varies and that recombination rate and mutation rate are correlated.

### Conclusions

The idea of widespread interbreeding between modern humans and other hominids has been broadly and rapidly accepted, I am sure in part because the idea of carrying their legacy is undeniably romantic. Another key element is that, so far, a plausible alternative hypothesis has not been available. Mounting evidence in favour of HI offers an alternative explanation for many or most of the observations used to infer introgression, from differential patterns of base sharing to changes in apparent block size. The HI-mediated mutation slowdown hypothesis fits better with a range of direct tests of fit in which opposing predictions of the two hypotheses are compared. Consequently, there is now a clear need to explain why mutation slowdown is so strongly favoured before the idea of a widespread Neanderthal legacy can be considered proven.

## Methods

### Data

Modern human sequences were obtained from the 1000 genomes project, Phase 3, and downloaded as composite vcf files. These comprise low coverage genome sequences for 2504 individuals drawn from 26 modern human populations spread across five main geographic regions: Africa (**LWK**, Luhya in Webuye, Kenya; **GWD**, Gambian in Western Division, The Gambia; **YRI**, Yoruba in Ibadan, Nigeria; **ESN**, Esan in Nigeria; **MSL**, Mende in Sierra Leone; ACB, African Caribbean in Barbados; ASW, African Ancestry in Southwest US), Europe (**GBR**, British from England and Scotland; **FIN**, Finnish in Finland; CEU, Utah Residents (CEPH) with Northern and Western Ancestry; **TSI**, Toscani in Italy; **IBS**, Iberian populations in Spain), Central Southern Asia (GIH, Gujarati Indian in Texas; **PJL**, Punjabi in Lahore, Pakistan; **BEB**, Bengali in Bangladesh; STU, Sri Lankan Tamil in the UK; ITU, Indian Telugu in the UK), East Asia (**CHS**, Han Chinese South; **JPT**, Japanese in Tokyo, Japan; **CDX**, Chinese Dai in Xishuangbanna, China; **KHV**, Kinh in Ho Chi Minh City, Vietnam; **CHB**, Han Chinese in Beijing, China) and the Americas (PUR, Puerto Rican from Puerto Rico; CLM, Colombian in Medellin, Colombia; MXL, Mexican ancestry in Los Angeles, California; PEL, Peruvian in Lima, Peru). Codes in bold are populations sampled from their geographic origin, assumed to be less admixed than those sampled from elsewhere, unbolded. Although in the 1000 genomes dataset singleton variants are probably correctly called, to guard against possible sequencing errors and to be maximally conservative these were excluded from all analyses. A list of informative bases for the Altai – chimpanzee (PanTroX) – Hg19 alignment was kindly provided by Andrea Manica and Marcos Llorente.

### Analyses

All analyses were conducted using custom scripts written in C++, available on request. Human – chimpanzee alignments were extracted from the Ensembl-Compara eight primate alignments (http://www.ensembl.org/info/data/ftp/). To avoid alignment ambiguities, bases were only accepted if they lay within blocks of at least 300 bases and were 10 or more bases from the nearest gap, even if this was a single base indel.

## Acknowledgements

I have had a number of useful and informative discussions Anders Eriksson, Andrea Manica, Mathias Meyer, Rob Foley, Marta Lahr, Eske Willerslev, Martin Sikora, Simon Martin and Mathias Currat. This work was not funded.

