## Supplementary Material for "The quantity of Neanderthal DNA in modern humans: a reanalysis relaxing the assumption of constant mutation rate" (from another version of this preprint: version 1, 2016-07-22)

### 1. The back-mutation rate required to explain a signal of introgression.

Ostensibly, the likely number of back-mutations required to account for the asymmetrical ABBA / BABA counts appears to be too low. Thus, Green et al. report 8156936 instances of BBBA (Table S1 below), where the chimpanzee differs from both the two humans and from the Neanderthal, consistent with the classic approximation of 1% divergence between humans and chimpanzees. Assume the mutation rate is $10^{-8}$ per base per generation [1], distributed randomly but evenly across the genome, and the time available for back-mutations to occur differentially between Africans and non-Africans is 50,000 years or 2,000 generations. Then, only 163.1 ABBA and BABAs would be formed, far fewer that the ABBA-BABA difference of around 8,000. Similarly, based on the same mutation rate and assuming a modern human common ancestor living 750,000 years or 30,000 generations ago, the number of back-mutations in a typical modern human would be 2,500. Again, this is far fewer than the observed number of ~200,000 ABBAs + BABAs together [2].

However, the above calculation is naïve, and the real number of back-mutations is likely to be substantially higher. This is shown empirically by the same data table in Green et al., who list just over 6000 instances each of BCBA and CBBA (Table S1 below). Both these base conformations require two independent mutations at the same site, at least one of which must be a transversion. Since transitions outnumber transversions by around two to one despite representing only half the possible changes, the relative probability of a transition to a transversion is around four [3], implying around 60,000 ABBAs and BABAs due to back-mutation in the Green et al. dataset and 240,000 genomewide. Moreover, even this number is likely to be an under-estimate. First, sites with three plus states are probably under-reported. Second, site-specific and sequence-context considerations [3 4 5] may cause some sites to mutate mainly by transitions while others would favour transversions. Third, in yeast, heterozygous sites are recognised and 'repaired' by gene conversion. With the same biochemical pathways in higher organisms and mounting evidence for heterozygote instability in humans, it is reasonable to assume that the same process operates. If so, then this would constitute a directed mechanism that actively promotes back-mutations.

Another approach to discover whether the rate of back-mutations is in the correct range to account for the ABBA-BABA asymmetry is to measure their frequency semi-directly. Consider two sites so close together that recombination can be ignored, say <200bp apart. This is consistent with most SNPs in the 1000 genomes Phase 3 data. Sites 1 and 2 initially carry ancestral alleles A1 and A2 respectively. Subsequently, each site mutates to a derived allele, D1 and D2. The result is two SNPs that between them carry three haplotypes, of which one haplotype is always the un-mutated A1A2. Depending on which chromosomes mutate there are three possible combinations for the two mutated haplotypes: A1D2 & D1D2; D1A2 & A1D2; and D1A2 & D1D2. In each case at least one 2-locus genotype is impossible, referred to henceforth as a discordant genotype. For example, the combination (A1A2, A1D2, D1D2) cannot produce individuals that are homozygous derived at the first SNP and homozygous ancestral at the second (i.e. D1D1 / A2A2). Ignoring recombination, discordant genotypes cannot arise without multiple mutations at one (or both) of the two sites.

**Table S1. Conformation counts for a French-San-Neanderthal-chimpanzee alignment, redrawn from Green et al. to emphasise pairs of conformations where only the two humans differ.** For the commonest four pairs of states (highlighted) asymmetrical counts are found, though for BCBA / CBBA the difference is non-significant. Note, the Table appears to contain a typo. The ABAA / BAAA pair is listed such that the commoner conformation is the one where 'B' occurs in the French. This is at odds both with the general observation that Africans carry more substitutions than non-Africans and the data on the previous page, p137 of their Supplementary Material, where the counts are given the other way round.

| F-S-N-C | Counts | Ratio |
|---|---:|---|
| AAAA | 818,322,920 | |
| AABA | 5,827,247 | |
| ABAA | 689,594 | |
| BAAA | 756,324 | 1.097 |
| ABBA | 95,347 | |
| BABA | 103,612 | 1.087 |
| ABCA | 2,995 | |
| BACA | 3,544 | 1.183 |
| BCBA | 6,147 | |
| CBBA | 6,264 | 1.019 |
| BBAA | 303340 | |
| BBBA | 8156936 | |
| BBCA | 32607 | |
| BCAA | 972 | |
| BCDA | 36 | |

To screen for discordant genotypes, I analysed all adjacent pairs of SNPs in each of the 26 populations in the 1000 genomes Phase 3 data [1]. For each SNP pair I used the observed two-locus genotype frequencies to calculate the best-fit frequencies of all four possible haplotypes (A1A2, A1D2, D1A2, D1D2) and also for each possible subset of three haplotypes. For both the four-haplotype and the best three-haplotype solutions I calculated a chi-squared goodness of fit and then sought loci where the best three-haplotype solution was either impossible (a genotype exists in multiple copies across the 1000g dataset despite an expected frequency = 0) or was significantly (chi-squared >10.827, P<0.001) less likely than the four-haplotype solution.

In total, this approach yields an estimate of 40,000 locus-pairs across the autosomes at which double mutations have occurred. Moreover, there are several reasons why this is a large under-estimate. Most importantly, many double mutations will fail to generate the 4$^{th}$ haplotype, many simply regenerating repeat copies of existing haplotypes. If the three haplotypes occur at equal frequency, only one third of the possible mutations would create the 4$^{th}$ haplotype, implying that the real number of back-mutations is at least 120,000. Second, even when all four haplotypes have been generated, subsequent drift may cause one or more to be lost. The exact probability of this depends on population size, population history and sampling strategy for the current data, but the number of sites lost in this way is likely to be appreciable. Third, the Phase 3 data are based on low-coverage genome sequences and include extensive interpolation to produce allele calls at all sites in all individuals. Since interpolation uses existing haplotypes to infer missing data, this process can never generate discordant haplotypes that do not exist elsewhere in the dataset. Thus, taken together, these approximate calculations support the previous analyses in suggesting that back-mutations are far commoner than naïve calculations suggest. Crucially, back-mutations appear common enough to provide a viable alternative hypothesis that is at least potentially capable of accounting for the observed ABBA-BABA signal.

**2. Dependence of back-mutation rate on outgroup distance.**

I am keen to distinguish between introgression and back-mutation as the two possible explanations of the observed asymmetry in ABBA and BABA counts. One prediction of the back-mutation hypothesis is that the ABBA-BABA signal will be an endogenous property of the two humans and therefore will persist regardless of whether the Neanderthal genome is present as a reference. I therefore analysed tetrads comprising Human1 – Human2 – AA – chimpanzee, where AA is the ancestral human allele inferred from a panel of higher primates as given in Phase 3 of the 1000 genomes project [1]. To prevent any possible spill-over signal, all informative Neanderthal sites were excluded. Just as with the Neanderthal genome, relative to Africans, Europeans exhibit excess base-sharing with the ancestral allele. However, the size of this excess is more than double that seen when the Neanderthal genome is used. The fact that the signal persists even when all Neanderthal information is removed shows that Neanderthal introgression is unlikely to be main underlying cause but raises questions about why the signal gets stronger.

A stronger signal is expected under the back-mutation model for at least two reasons. First, the inferred ancestral allele by definition lies at the base of the human clade, so forms a shallower three-way clade that the Human1 – Human2 – Neanderthal trio. Consequently, the number of recently derived mutations will be reduced and those that occurred after the 'out of Africa' event will make up a relatively higher proportion. Second, mutation rate not only varies greatly across the genome and between sites, with some bases having much higher mutation probabilities than others [3], but also likely varies through time. This can be inferred from two observations: first, that mutation rate and recombination rate are correlated [6]; and, second, that recombination hotspots differ considerably between humans and chimpanzees [7]. By implication, mutation hotspots also vary in genomic location over short evolutionary timescales [8]. If so, then the shallower time-depth of the three-way clade, the greater the number of hotspots that will have remained in the same location over the time considered. In turn, this should increase the probability of a back-mutation.

As a further test I also conducted ABBA-BABA tests for various combinations of higher primates. Asymmetrical ABBA-BABA counts are also found for Human1 – Human2 – chimpanzee – gorilla and for Human1 – Human2 – chimpanzee – orangutan. Here the signal, although highly significant, is appreciably weaker than for either the Neanderthal or the ancestral allele analyses. Inter-breeding between chimpanzees and modern humans is biologically implausible. However, a weak ABBA-BABA signal is consistent with mutation slowdown because under this hypothesis the ABBA-BABA signal is endogenous to the Human1-Human2 pair and should be present but progressively weaker with increasing distance between the test group and the common ancestor of modern humans.

**3. Recent derived alleles are commoner in Africa.**

The mutation slowdown hypothesis predicts that mutations that occurred since humans migrated out of Africa will be less common outside Africa due to the loss of heterozygosity causing a parallel reduction in mutation rate. Data are not available to estimate the mutation rate directly in different human populations. However, to a close approximation, rare alleles are almost invariably recent such that alleles present in only two copies in the 1000 genomes Phase 3 data are estimated to be on average around 150 generations old [9]. I therefore took 10% as the threshold likely below which most alleles are likely to be descendants of mutations that occurred since humans left Africa ~2,000 generations ago. I refer to these as recent derived alleles and counted them in each of the 26 populations represented in 1000g. To reduce biases that may accrue due to differences in population size and relatedness within samples I defined allele frequency as the average frequency across all populations rather than the frequency in any particular population.

Counts for recent derived alleles are presented in Figure S1. Typical frequencies are 3.37 fold higher in African compared with non-African populations. Moreover, the strongly admixed American populations show distinctly higher counts, consistent with European admixture enhancing local heterozygosity, thereby increasing (slightly) the mutation rate and hence the number of recent derived alleles. Conversely, the two righthandmost African populations show somewhat lower counts, consistent with these being the somewhat admixed Afro-Caribbean and African Americans sampled from Texas. Here, low levels of European ancestry will likely reduce the number of recent derived alleles by more than would be generated by any slight increase in heterozygosity.

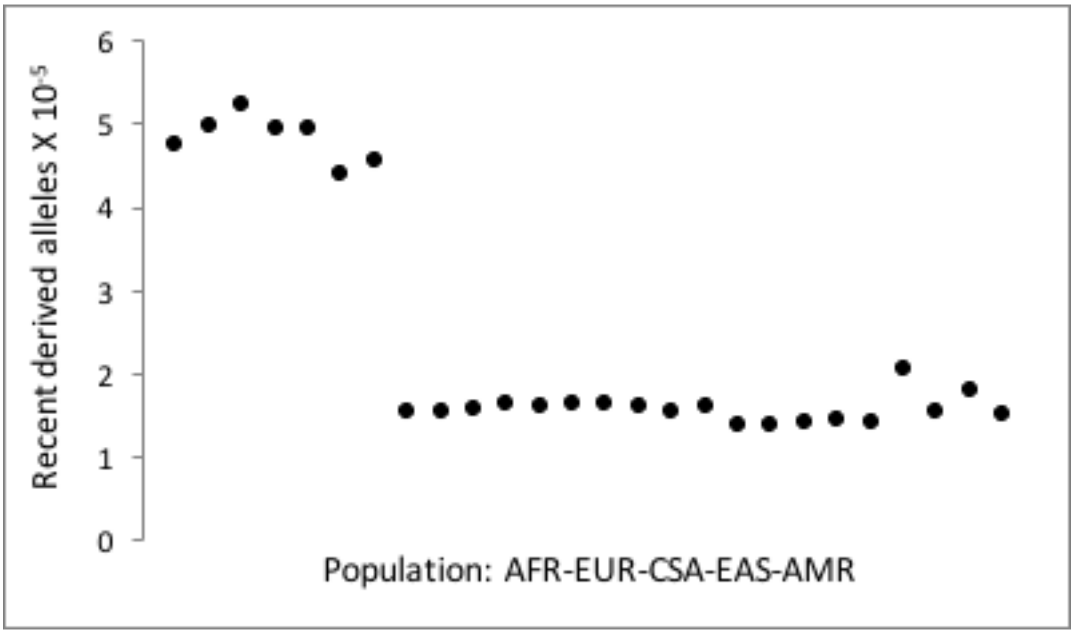


**Figure S1. Recent derived alleles in diverse human populations.** Recent derived alleles are defined as non-ancestral alleles with an average frequency across all 26 1000 genome Phase 3 populations of under 10%. Populations are arranged by major population group: 7 x African (AFR), 5 x European (EUR), 5 x Central Southern Asian (CSA), 5 x East Asian (EAS) and 4 x American (AMR). Populations appear in the same order as given in Methods. The Y-axis is the sum of the frequencies of qualifying alleles in each population and hence is the expectation for the number of recent derived alleles in a typical individual from that population, divided by $10^5$.

**4. Searching for unusual fragments outside Africa.**

To test whether non-Africans carry unusual fragments consistent with a Neanderthal legacy I used the 1000 genomes data and calculated the maximum within-population pairwise divergence (MWD) in each population based on non-overlapping 20Kb windows. Neanderthal sequences in non-Africans should cause the windows in which they fall often to show MWD values that exceed the equivalent value in Africa, where introgression is assumed to be rare or absent [2]. Windows typically carry of the order of 50 plus base differences between a random pair of individuals. To minimise stochastic noise from partial windows and from relatively monomorphic regions, I excluded all windows where the mean inter-individual difference in Africa fell below <15 (approximately two standard deviations below the mean). This left 127,370 informative autosomal windows equivalent to 2.34Gb.

Reliable values for the expected excess level of Neanderthal divergence are difficult to find. Data in Green et al. (see Table S1 above) suggest a value of 4.2X based on base combination counting. However, a more realistic / conservative value is 1.5X, measured from the autosomal tree in Figure 2 of Prufer et al. [10]. Turning to the 1000 genomes data, only 92 windows (0.072%) yield values where MWD in any of the European populations exceeds 1.5X the MWD in Africa.

It could be argued that the reason a low proportion of windows appear to carry unusual fragments is that selection has acted to remove more divergent fragments or, perhaps, that mean fragment size is smaller than expected, such that most introgressed fragments are much smaller than the windows in which they appear. To address these possibilities, I compared the distribution of MWD values between the autosomes, where introgression has been inferred to be widespread, and the X chromosome, where little or no evidence has been found. If windows where MWD in Europe is higher than MWD in Africa genuinely reflect genomic regions where introgressed fragments persist, such windows should be appreciably rarer and / or smaller on the X chromosome. In fact, MWD in Europe and Asia exceeds MWD in Africa in 11 of 7462 informative windows (0.14%) on the X, approximately double the average number on the autosomes.

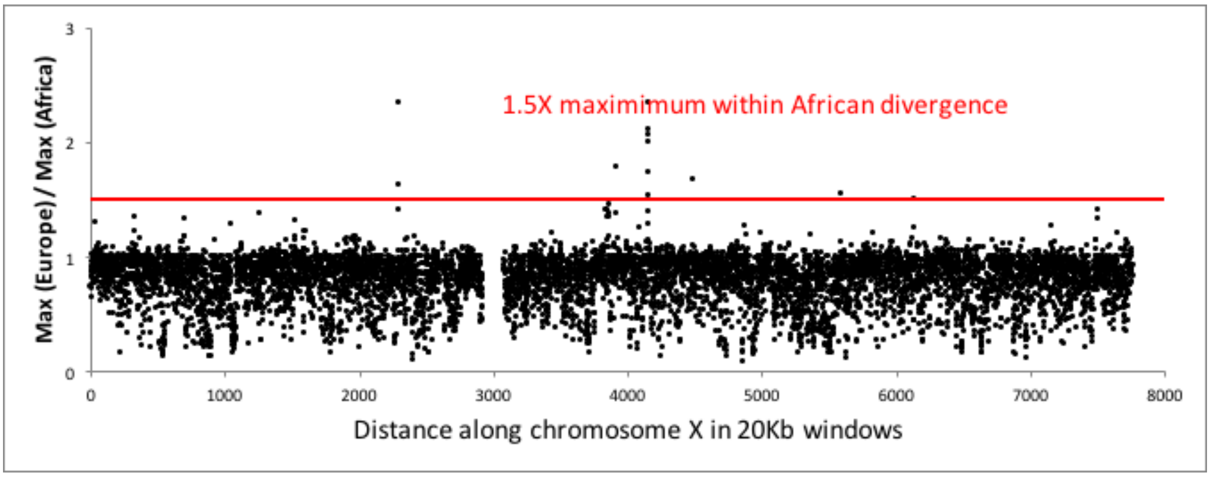


**Figure S2. A test for unusual fragments on the X chromosome in non-Africans.** Data from the 1000 genomes Phase 3 data were partitioned into non-overlapping 20Kb windows. Within each window and each of the 26 populations all-against-all divergences were calculated and the maximum value recorded for each population. In the graph, the Y-axis is the ratio of the largest maximum found outside Africa (here taken as Europe + East Asia, the regions where most Neanderthal introgression has been inferred) to the largest maximum found within Africa (LWK, GWD, ESN, MSL & YRI). A Neanderthal fragment outside Africa might be expected, on average, to generate a ratio of approximately 1.5X, as indicated, or more conservatively, greater than unity. In practice, the vast majority of windows yield ratios less than unity. Importantly, the frequency and magnitude of high ratios on the X is similar to and if anything greater than is seen on the autosomes, suggesting strongly that these peaks are largely not related to introgression.

A further possible issue relates to the fact that genetic divergence accrues stochastically and varies between genomic regions. Consequently, the modest average excess Neanderthal-human divergence (relative to human-human) may be localised in a rather small proportion of all windows, the rest failing to stand out as being unusually discordant.  To estimate the size of this problem I ‘spiked’ each non-African population with a single, randomly-selected African individual from population GWD. I then recorded the number of windows where the pairwise comparison that contributed the MWD in Europe / Asia involved the introduced African individual.  Since Neanderthals should be even more dissimilar, the number of windows found, 33%, sets a minimum for the proportion of all windows in which a Neanderthal fragment would be detected.  From this, it seems reasonable to assume that upwards 50% of all windows would show sufficient Neanderthal-human divergence for a Neanderthal fragment to stand out as being unusual in Europe.


1 1000 Genomes Project Consortium. A map of human genome variation from population-scale sequencing. *Nature* **467**, 1061-1073 (2010).

2 Green, R. E. *et al.* A draft sequence of the Neandertal genome. *Science* **328**, 710 (2010).

3 Ségurel, L., Wyman, M. J. & Przeworski, M. Determinants of Mutation Rate Variation in the Human Germline. *Ann. Rev. Genomics Hum. Genet.* **15**, 47-70, doi:10.1146/annurev-genom-031714-125740 (2014).

4 Zhang, W., Bouffard, G. G., Wallace, S. S. & Bond, J. P. Estimation of DNA sequence context-dependent mutation rates using primate genomic sequences. *J. Mol. Evol.* **65**, 207-214 (2007).

5 Vowles, E. J. & Amos, W. Evidence for widespread convergent evolution around human microsatellites. *PLoS Biol.* **2**, e199 (2004).

6 Duret, L. & Arndt, P. F. The impact of recombination on nucleotide substitutions in the human genome. *PLoS Genet.* **4**, e10000071 (2008).

7 Ptak, S. *et al.* Fine-scale recombination patterns differ between chimpanzees and humans. *Nat. Genet.* **37**, 429-434 (2005).

8 Amos, W. Even small SNP clusters are non-randomly distributed: is this evidence of mutational non-independence? *Proc. R. Soc. B* **277**, 1443-1449 (2010).

9 Mathieson, I. & McVean, G. Demography and the Age of Rare Variants. *Plos Genetics* **10**, doi:10.1371/journal.pgen.1004528 (2014).

10 Prufer, K. *et al.* The complete genome sequence of a Neanderthal from the Altai Mountains. *Nature* **505**, 43-+, doi:10.1038/nature12886 (2014).
